# Engineered Protein Nanosheets for the Scale up of Mesenchymal Stem Cell Culture on Bioemulsions

**DOI:** 10.1101/2023.12.18.572135

**Authors:** Minerva Bosch-Fortea, Clemence Nadal, Alexandra Chrysanthou, Daniele Marciano, Hassan Kanso, Nam Nguyen, Julien E. Gautrot

## Abstract

The rapid progress in cell therapies and stem cell technologies requires the development of novel bioprocessing and biomanufacturing pipelines able to cope with the scale up of cell manufacturing. In this respect, microdroplet technologies have already revolutionised the field of biotechnologies, but remain ill-suited to the culture of adherent cells. In this report, we describe the engineering of albumins with cell adhesive peptides for the stabilisation of microdroplets enabling the scale up of mesenchymal stem cell (MSC) expansion. We characterise the modified albumins prior to study their self-assembly at liquid-liquid interfaces via interfacial shear rheology, and mechanical strengthening through the formation of crosslinked nanosheets. The biofunctionalisation of these protein nanosheets is then characterised by fluorescence microscopy. In turn, the ability of the resulting bioactive microdroplets to promote rapid cell adhesion and expansion is examined and the extensive deposition of matrix associated with such cultures is characterised. The culture of MSCs is then scaled up 100 fold, first at the surface of fluorinated oil emulsions, then on plant-based emulsions stabilised by engineered protein nanosheets and the phenotype of resulting cells is characterised. The microdroplet culture system presented displays attractive advantages over existing technologies, in terms of simplicity of processing, compatibility with regulatory expectations and costs of production, and offers exciting opportunities for translation to cell manufacturing, for cell therapies and cultivated meat applications.

## Introduction

In the last three decades, progress in the field of stem cell biology and regenerative medicine has enabled the demonstration of a range of different cell therapies enabling to tackle a wide range of diseases and conditions, from wound healing and bone reconstruction to cardiac repair, ocular regeneration and lung repair [1–5]. Similarly, cell-based therapies have been proposed to correct deficiencies in the production of factors and hormones, such as Factor VIII and insulin [6–9], or the production of exosomes [10–12]. Although these studies have demonstrated promising performance in small and sometimes large animals, their translation to the clinic has been slow, owing to a range of different factors, including poor translation to a human context and human biology, poor cell retention and viability, an immune response, the difficulty to deliver cells using scaffolds approved by regulators and the manufacturing of sufficient cell densities for implantation [13–15]. Indeed, although cell densities of 1-5 million per animal are often sufficient to demonstrate efficacy in mouse or rat models, translation to larger animals and patients would typically require the implantation of several hundred million cells. The scale up of cell manufacturing therefore remains an important hurdle to address to enable the translation of cell therapies to the clinic. Similarly, the scale up of biotherapeutics production, whether defined factors or exosomes, will require novel platforms to reduce costs and limit the potential for contamination of products [14]. These requirements are also shared by emerging applications of cell-based technologies, such as cultivated meat production, which remains prohibitively costly and difficult to scale up to the potential consumer demand [16,17].

A number of key challenges remain in order to address pitfalls associated with current cell manufacturing. Although some mammalian cell lines can be engineered or selected for culture in suspension, many cell lines studied at the laboratory scale and proposed for translation (apart from CAR-T cells and pancreatic islet cells, for example) are adherent and require solid substrates or hydrogels to enable adhesion and spreading, as well as expansion and the retention of cell phenotypes [18–20]. The gold standards in this area consist in cultivating adherent cells, such as mesenchymal stem cells or fibroblasts, on solid microcarriers (typically polystyrene modified with cell adhesive peptides), which can be scaled up in stirred tank bioreactors [21,22]. However, recovery of the resulting cell products and separation from microcarriers in mild conditions is difficult to scale up and limits translation. In addition, microcarrier solutions remain expensive (estimated between 40 and 80 % of the total cost of cell culture reagents; based on the cost of Synthemax microcarriers for the culture of a range of different adherent cell types) and lead to contamination of cell products and biotherapeutics with microplastics.

Recently, liquid substrates, in the form of oil microdroplets, were proposed to replace solid microcarriers. This has enabled the culture and retention of phenotype of a broad range of cell types, from fibroblasts, keratinocytes and HEK293 cells, to mesenchymal stem cells (MSCs), induced pluripotent stem cells and neural cells [23–28]. Underpinning these technologies, it was demonstrated that protein nanosheets self-assembling at corresponding liquid-liquid interfaces and presenting strong interfacial mechanical properties enabled the adhesion, spreading and proliferation of cells [28–32]. In order to support cell adhesion and expansion at the surface of oil microdroplets, protein nanosheets were proposed to require a combination of tensioactive (stabilising droplets), scaffolding (strengthening interfacial mechanics to resist cell adhesion) and bioactive properties (engaging with cell membrane receptors such as integrins to sustain cell adhesion). The resulting bioactive emulsions, or bioemulsions [25,26], have been shown to enable the retention of stem cell phenotype [24,26,33,34], even over prolonged periods of time corresponding to multiple passages [33], to support the secretion of biotherapeutics including exosomes [35], and to enable the formation of biomimetic niches for haematopoietic stem cells [34]. However, most of these systems relied on fluorinated oils that do not enable direct implantation, and likely will require separation from final cell products. In addition, the proteins used required co-surfactants such as pentafluorobenzoyl chloride or perfluorooctanoyl chloride, which should be replaced or avoided prior to translation. The extra- cellular matrix protein fibronectin, enabling integrin-mediated cell adhesion, was showed to enable direct nanosheet formation without additional of additional co-surfactant or crosslinker [27], likely due to its ability to fibrillate at some hydrophobic interfaces [36,37], but is poorly tensioactive. On the other end, β-lactoglobulin, a protein excellent at stabilising emulsions and displaying inherent scaffolding properties, required biofunctionalisation with cell adhesive peptides [25].

In this report, we engineer bovine serum albumin (BSA) with functional groups enabling the coupling of cell adhesive peptides via radical thiol-ene coupling and Michael addition (Figure 1). The resulting proteins are then used to stabilised oil microdroplets and further crosslinked to confer scaffolding properties, in a scalable format. We investigate the interfacial shear mechanical properties of corresponding interfaces during these processes and characterise the stability of associated emulsions. We then compare the ability of these different proteins to promote the adhesion and expansion of bone marrow-derived MSCs at the surface of microdroplets, first in the context of fluorinated oil, then plant-based (rapeseed oil) interfaces. We further demonstrate the scale up of cell culture on these bioemulsions, in conical flask bioreactors, to scales relevant to pre-clinical studies (100 fold). Finally, we characterise the retention of stem cell marker expression upon culture on corresponding bioemulsions and demonstrate the simple recovery of cells from these culture systems via a range of strategies.

**Figure 1.**
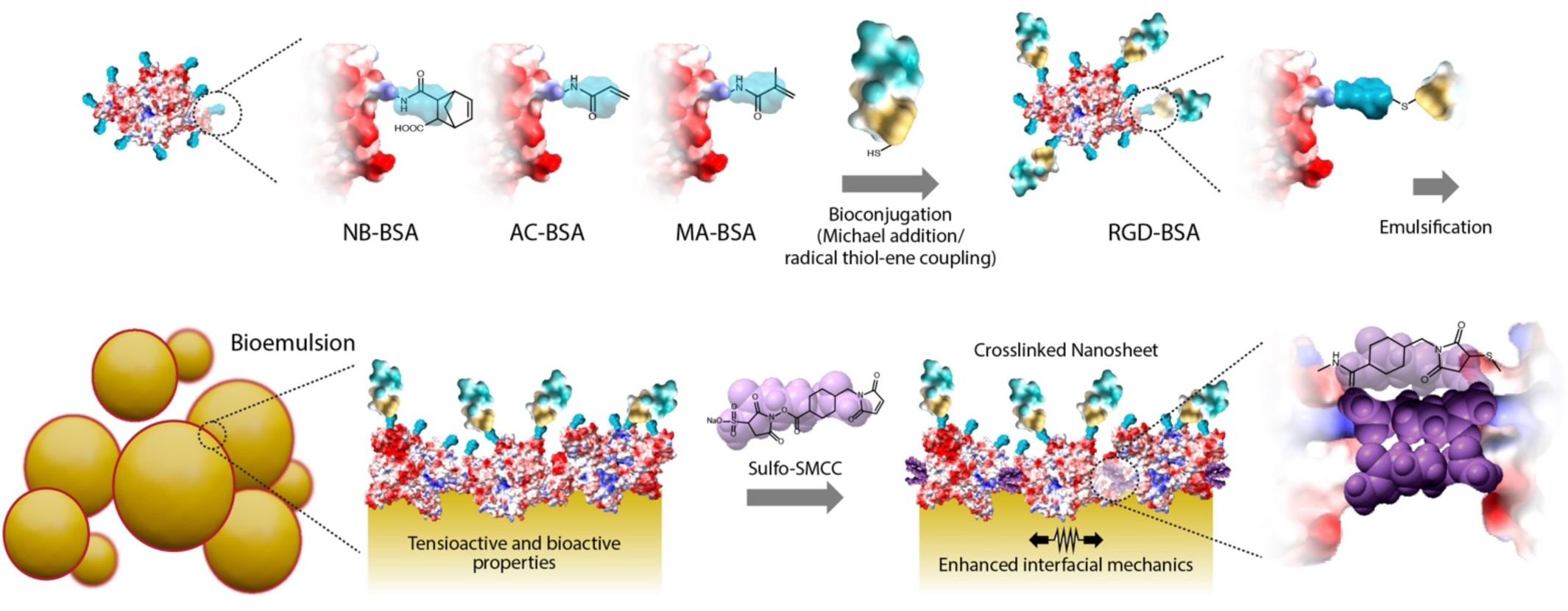
Schematic representation of protein nanosheets stabilising microdroplets, functionalised with norbornene (NB-BSA), acrylate (AC-BSA) and methacrylate moieties (MA-BSA). Bioconjugation with RGD peptides via Michael addition or radical thiol-ene coupling is followed by emulsification and sub-sequent further crosslinking with sulfo-SMCC.

## Materials and Methods

### Synthesis of functionalized BSA

NB-BSA was obtained using the following protocol. 10 g of BSA (15 mmol, Sigma-Aldrich) were dissolved in 100 mL of carbonate (NaHCO3/Na2CO3) buffer at pH 9. Then, 2 g of carbic anhydride (12.2 mmol, Thermo Fisher Scientific) were slowly introduced into the reaction mixture, at room temperature. The resulting mixture was then heated whilst stirring at 40 °C for 4 h. A pale yellow solution was recovered, dialysed against deionised water for 4 days and freeze-dried. The functionalisation degree of the resulting NB-BSA was determined by ^1^H NMR, by integration of the 2 alkene protons (δ 6.0- 6.3 ppm) of norbornene moieties, normalised by the aromatic protons of BSA (δ 6.5-7.5 ppm).

### RGD coupling to biofunctionalised BSA

RGD coupling to functionalized BSA molecules was performed prior to formation of bioemulsions. A ration of RGD peptide to functionalised BSA molecules of 10 was targeted. Peptides were selected to enable enhanced reaction with functional groups in the modified BSA backbone. For NB-BSA, a GCGGRGDSPG peptide was selected for coupling via radical thiol-ene chemistry, using a solution of 0.03 wt % lithium phenyl-2,4,6-trimethylbenzoylphosphinate (1 mM; LAP, Sigma-Aldrich), 0.1 mg/mL of peptide and 1 mg/mL of BSA, under irradiation with 405 nm blue light for 3 min (at 20 mW/cm^2^). For AC-BSA and MA-BSA, a CGGRGDSPG peptide was conjugated via Michael addition in PBS pH 9 for least 3 h (10 mL of buffer, 0.1 mg/mL of peptide and 1 mg/mL of BSA). The same procedure was used for CGGRGDSP*K*-FITC-G and CGGRGDSP*K*-FITC-G (FITC bearing) peptides, for AC-BSA/MA-BSA and NB-BSA, respectively.

### Preparation of bioemulsions

Emulsions were generated by mixing fluorinated oil (Novec 7500, ACOTA) and Rapeseed oil (Sainsbury’s) with the protein aqueous solution (1 mg/mL), at a 1:2 volume ratio. The vial was shaken for 15 s using a vortex and incubated for 1 h at room temperature. The aqueous phase was aspirated, and the emulsions thoroughly washed with PBS to remove excess protein. For those bioemulsions that were crosslinked, sulfo-succinimidyl 4- (N-maleimidomethyl) cyclohexane-1-carboxylate (sulfo-SMCC, Thermofischer Scientific) was dissolved (2 mg/ mL) into 0.5 mL of distilled water and sonicated for two minutes. Bioemulsions were then incubated in this solution for 1 h at RT. The aqueous phase was aspirated and replaced with PBS six times to remove the excess sulfo-SMCC. For cell culture, bioemulsions were transferred to either 24-well plates or conical flasks with vented caps treated with poly(L-lysine)-graft-polyethylene glycol (PLL-g-PEG, SuSoS; 25 μg/mL) for 1 h. *^1^H NMR.* ^1^H NMR spectroscopy was carried out at 298 K using a Bruker AVIII 400 instrument. Chemical shifts δ are referenced to the residual solvent peak (D2O : H = 4.79 ppm).

### Circular Dichroism

Protein solutions (BSA, NB-BSA, AC-BSA and MA-BSA) were prepared in deionized water at a final concentration of 1 mg/mL. The samples were kept at room temperature and used for measurements within 2 hours. 350 μL of each solution was transferred to a sample cuvette of 1 mm thickness and measurements were carried out at 25°C in a Chirascan V100 CD spectrometer. Data analysis was performed smoothing measurements using the Savitzky-Golay smooth filter with a script written in MATLAB.

### Gel Electrophoresis

Samples were mixed with 2 x SDS, boiled at 95°C for five min then run on a 4-20% SDS page gel with Tris-glycine SDS running buffer at 230 V for 30 min. Gels were stained with InstantBlue® Coomassie Protein Stain solution for 1 h and imaged under a transilluminator.

### Mastersizing and Analysis of Bioemulsion Stability

The stability of bioemulsion was evaluated at various timepoints using a Mastersizer 2000. Particle size distribution was analysed for each condition on the day of bioemulsion formation (Day 1), as well as after 7 and 15 days of storage at 37 °C with continuous agitation with an orbital shaker (60 rpm). Additionally, bright-field microscopy was employed to monitor emulsion stability.

### Interfacial Shear Rheology Measurements

Interfacial shear rheological measurements were carried out on a Discovery Hydrid-Rheometer (DHR-3) from TA Instruments, using a Du Nouy ring geometry and a Delrin trough with a circular channel. The Du Nouy ring has a diamond-shaped cross section that improves positioning at the interface between two liquids to measure interfacial rheological properties while minimizing viscous drag from upper and sub-phases. The ring has a radius of 10 mm and is made of a platinum−iridium wire of 400 μm thickness. The Derlin trough was filled with 4 mL of fluorinated oil (Novec 7500, ACOTA). Using axial force monitoring, the ring was positioned at the surface of the fluorinated oil and was then lowered by a further 200 μm to position the medial plane of the ring at the fluorinated phase interface. 4 mL of the PBS solution was then gently introduced to fully cover the fluorinated sub-phase. Time sweeps were performed at a frequency of 0.1 Hz and temperature of 25 °C, with a displacement of 1.0 × 10^−3^ rad (strain of 1%) to follow the self-assembly of the proteins at corresponding interfaces. 15 min after the beginning of the time sweep, the protein solution (100 mg/mL in PBS) was added in the aqueous phase to a final concentration of 1 mg/mL. In the case of non-crosslinked albumins interfaces, the time sweep was carried out for 3 additional hours.

In the case of crosslinked albumin nanosheets, the remaining soluble proteins were removed after 1 h by washing the aqueous phase for 30 min with a microfluidic flow controller (OB1 MK4 pressure controller ElveFlow, Elvesys, France), which allowed the regulation of inlet and outlet PBS flow rates (overall 1 mL/min). Following this washing cycle, a sulfo-SMCC solution was injected in the aqueous phase to a final concentration of 2 mg/mL. The excess crosslinker was removed by a 30 min washing cycle after 1 h.

In both crosslinked and non-crosslinked systems, frequency sweeps (with displacements of 1.0 × 10^−3^ rad) and amplitude sweeps (at a frequency of 0.1 Hz) were carried out at 25 °C to examine the frequency-dependent behaviour of corresponding interfaces and to ensure that the selected displacements and frequencies were within the linear viscoelastic region.

Stress relaxation was performed successively at 0%, 0.1 %, 0.5 % and 1% strain for 120 s. Stress relaxation data from the 10^th^ second onwards, when the constant strain was maintained, was plotted in OriginPro, and fitted with double exponential decay fit, according to the following equation:

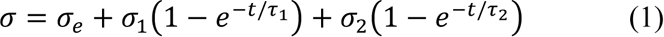

In this equation, σ is the measured residual stress, σe is the elastic stress and σ1 and σ2 are viscous relaxation components. The degree of stress retention (σr) is calculated as:

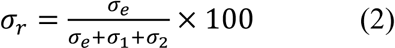

### Evaluation of emulsion stability

The stability of the different formulated bioemulsions was assessed at different timepoints using a Mastersizer 2000 according to manufacturer’s instructions. Particle size distribution was determined for each condition at the day of bioemulsion formation (Day 1) and after 7 days and 15 days kept at 37 °C under agitation in an orbital shaker (60 rpm). The emulsion stability was also monitored via bright-field microscopy.

### Mesenchymal stem cells culture and seeding on emulsions

Bone marrow-derived human mesenchymal stem cells (MSCs) were purchased from PromoCell and cultured in MSC Growth Medium 2 (PromoCell). MSCs between passages 2 and 6 were used for the experiments reported. For cell seeding on bioemulsions, MSCs were harvested using Accutase (PromoCell) and centrifuged at (200 x g) for 5 min. 3 10^4^ cells were seeded in 24-well plates containing 50 µL of bioemulsion and 750 μL of MSC Growth Medium and cultured for 7 days at 37 °C in a humidified atmosphere with 5% CO2. For the 2D controls, 7,000 cells/well were seeded in 24- well plates containing 500 μL of MSC Growth Medium. Half of the medium was replaced with fresh medium every two days.

### Culture in conical flask bioreactors

Scale up of the MA-BSA bioemulsions system was performed in 250 mL polycarbonate Erlenmeyer flasks (Corning) with Vent caps, using a total amount of medium of 50 ml. Volumes and amounts of further reagents and cells used where increased 100 times with respect to those used in 24 well-plate cultures. 5 ml of emulsions were used per conical flask. Cell densities (per surface area) used in bioemulsions and 2D conditions were 1600 cells/cm^2^, and matched based on an average diameter of 160 μm for the bioemulsions. Cells were kept under gentle agitation of 60 rpm on an orbital shaker placed inside an incubator at 37 °C in a humidified atmosphere with 5% CO2.

### Immunofluorescence staining

Cells cultured on 2D substrates (borosilicate coverslips) and on bioemulsions were fixed with 4% PFA for 20 min and then permeabilised with 0.2 % Triton X-100 for 10 min at 4 °C. Samples were then blocked with 3 wt% BSA at room temperature for 30 min. Primary antibodies were diluted in 3 wt% BSA and incubated from 1 h at 37°C, or overnight at 4°C at the recommended concentration, depending on the antibody. Primary antibodies and dilutions used were: anti-vinculin (V9264, Sigma-Aldrich, 1:500) and anti-fibronectin (F3648, Sigma-Aldrich, 1:500). After washing, secondary antibodies (Alexa Fluor 488/647-conjugated, Thermofisher, 1:1000), DAPI (Life Technologies, 1:1000) and phalloidin-TRITC (Sigma-Aldrich, 1:500) were incubated for 1 hour in the dark and then washed with PBS. Cells cultured as 2D monolayers were mounted with fluoromount (Thermo Fisher Scientific) over glass slides. Cells cultivated on emulsions were transferred to μ-slide 8 wells plates (Ibidi) and preserved with PBS+0.05% Azide (Sigma-Aldrich) at 4 °C for a maximum period of 2 weeks.

### Microscopy

A Zeiss LSM710 laser scanning microscope and a Nikon CSU-W1 spinning disk microscope with Two Photometrics Prime BSI sCMOS cameras were used for confocal imaging. Objectives used were usually EC Plan-Neofluar20x/0.5 M27 and Plan-Apochromat 63x/1.4 Oil DIC M27 on the Zeiss LSM710. Objectives used in the Nikon CSU-W1 were CFI Plan Apochromat VC 20X Air and CFI Apochromat TIRF 60XC Oil. Widefield fluorescent images were taken using a Leica DMi8 fluorescence microscope coupled to a Leica DFC9000 GT sCMOS camera. Objectives used were HC PL FLUOTAR L 20x/0.40 CORR PH1 and HC PL FLUOTAR L 40x/0.60 CORR PH2. Image analyses, z-stack projections, and 3D deconvolutions were carried out using ImageJ.

### Viability staining and cell counting

Cell viability was assessed using Live/Dead Viability/Cytotoxicity Kit (Invitrogen). A 300 mL sample of MSC-cultured emulsion was taken from the Erlenmeyer flask at different timepoints and placed in a 24-well containing a pre-warmed solution of PBS with Hoechst (1 mg/ml Thermofisher Scientific) and the Live/Dead staining solutions. After 30 min incubation, cells were imaged using a Leica DMi8 epifluorescence microscope. Viability was compared to that of cells growing on 2D tissue culture plastic at the same timepoints.

### Flow cytometry

Single cell suspensions of cells growing on bioemulsions and on 2D culture plastic were obtained via Accutase treatment and gentle agitation. Cells were then washed in 0.1% BSA and centrifuged (300 x g) for 10 min at 4°C. Cells were stained for 45 min at RT using fluorescence-labelled antibodies and Calcein Violet (#C34858, Thermo Fisher Scientific) for cell viability. Labelled cells were then washed in PBS + 1% BSA and analyzed on a FACS Canto II flow cytometer (BD Biosciences). Data analysis was performed using FlowJo software (Tree Star). Conjugated antibodies used were: FITC anti-CD73 (MCA6068F, Bio-Rad), APC anti-CD90 (MCA90APC, Bio-Rad), and PerCP/Cy5 anti-CD105 (323216, BioLegend).

### Re-seeding experiments

Cells grown on bioemulsions were reseeded to evaluate their capacity to adhere to other substrates after growing in these conditions. MSCs grown on rapeseed oil MA-BSA bioemulsions for 7 days in a 24-well plate were detached using Accutase for 5 min at 37 °C, reseeded on a glass coverslip and grown for 2 days. MSCs grown for 15 days on MA- BSA fluorinated oil emulsion in an Erlenmeyer flask were reseeded either on glass coverslips or in 10 mg/ml bovine fibrinogen (Sigma-Aldrich) and 2U/mL thrombin (Sigma-Aldrich) gels, after dissociation using different techniques. Cells were either seeded with the emulsions without detachment, after gentle disaggregation by pipetting up and down with a needle, and treating with Accutase for 5 min at 37°C.

### Statistical analyses

All statistical analysis was carried out using a GraphPad Prism 9 software. Two-tailed Student’s t-test or one way ANOVA with Tukey test for posthoc analysis were used to calculate statistical significance. The significance level was set at 0.05 and p-values are specified in the figure captions. Results are presented as the mean ± standard deviation (SD). Sample sizes and numbers of repeats are reported for each experiment in the figure captions.

## Results and Discussion

To develop a versatile modular platform enabling the formulation of bioemulsions, we selected albumins as globular proteins displaying inherent tensioactive properties. To introduce reactive moieties, acrylate, methacrylate and norbornene residues were coupled to bovine serum albumin, using corresponding anhydrides (Figure 1). This coupling step was carried out in carbonate buffers typically used for the functionalisation of amines with succinimidyl esters and anhydrides [38,39]. Following purification via dialysis, the resulting functionalised proteins were characterised via ^1^H NMR (Figure 2A). Analysis of the integration of the peaks in the olefin range (norbornene protons at δ: 6.0–6.3 ppm; acrylate protons at δ: 5.8–6.5; methacrylate protons at δ: 6.1–5.7 ppm), allowed the quantification of functionalisation levels, through comparison with aromatic protons of BSA near δ 7.5 ppm. Hence, we determined that 24 norbornyl, 71 acrylate and 28 methacrylate residues functionalised the corresponding NB-, AC- and MA-BSA proteins. These functionalisation levels were considered to be sufficient to enable effective bioconjugation of multiple peptides per protein.

**Figure 2.**
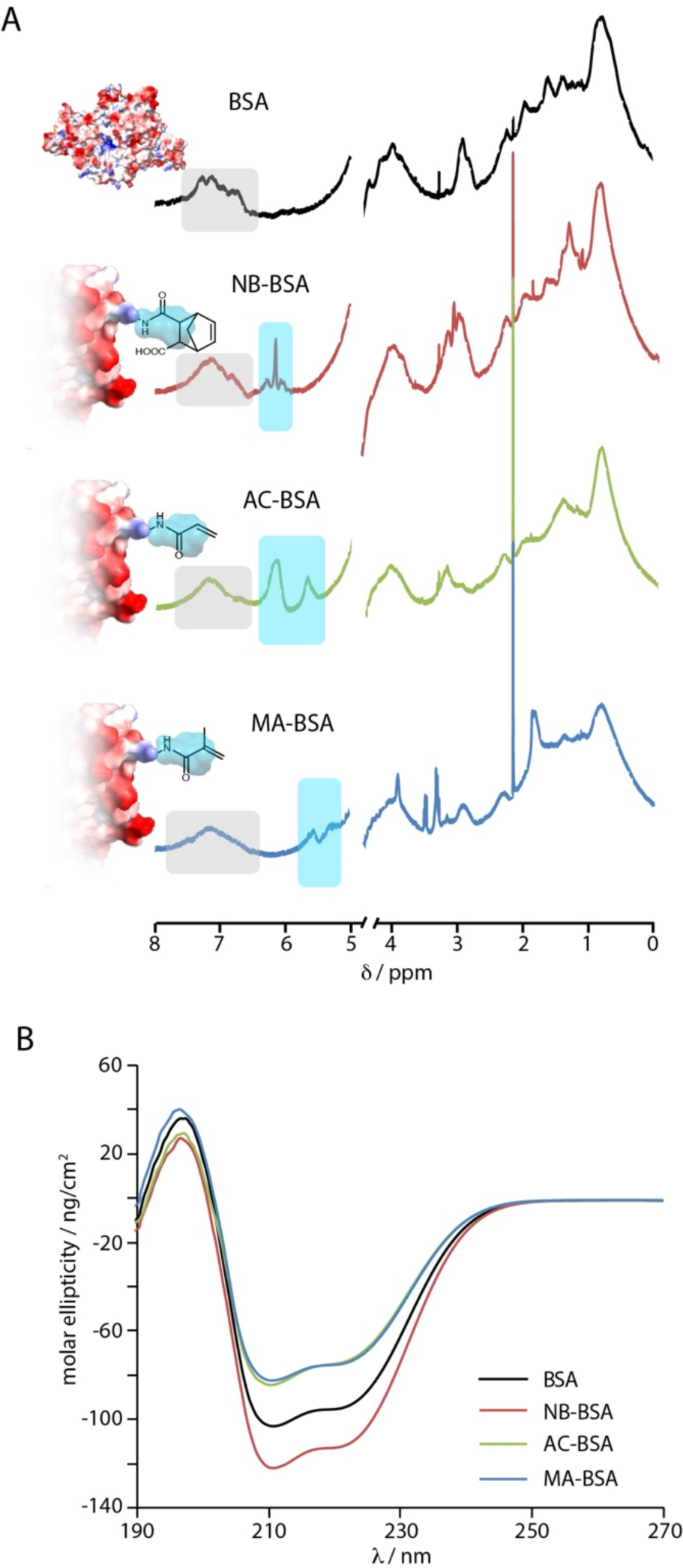
A. ^1^H NMR (D2O) spectra of BSA and functionalised BSA. The broken axis correspond to the position of H2O. Grey shaded areas correspond to protons from aromatic amino acid residues. The turquoise shaded areas correspond to the olefinic protons from norbornyl, acrylate and methacrylate residues. B. Circular dichroism spectra of modified BSAs compared to that of native BSA (BSA, NB- BSA, AC-BSA and MA-BSA were dissolved in PBS at a concentration of 1 mg/mL).

The impact of functionalisation on the structure of the resulting proteins was next assessed, as the globular conformation of albumins is considered essential to their tensioactive and self-assembly properties at hydrophobic interfaces [40,41]. Indeed, BSA is composed of multiple α-helices bundled together and forming a hydrophobic pocket enabling the binding of various apolar compounds, for example to modulate their transport [42,43], and underpinning assembly at hydrophobic surfaces [44,45]. Evaluation of protein structures through circular dichroism (CD) indicated the retention of highly α-helical architectures upon functionalisation (Figure 2B). Indeed, NB-BSA, AC-BSA and MA-BSA molecules displayed CD spectra presenting a characteristic positive peak at 196 nm and two negative peaks at 209 nm and 222 nm, comparable to those observed for pristine BSA and matching literature reports [46,47]. Only a moderate shift in the position of the 196 nm peak, and therefore loss of some of the α-helicity of the corresponding proteins, was observed. This retention in overall globular conformation is an excellent indicator that functionalisation levels should not interfere with self-assembly properties and the ability of corresponding albumins to stabilise emulsions. A globular architecture and striking changes in secondary structure were indeed associated with assembly at liquid-liquid interfaces, including hydrophobic and fluorophilic oil interfaces, and the stabilisation of associated emulsions [25,46,47].

The assembly of functionalised proteins at the surface of the fluorinated oil Novec 7500 was next examined, using interfacial rheology. This oil, often used in microdroplet microfluidic technologies [48,49], was selected as displaying high cytocompatibility, high density and low viscosity. It was found to support the adhesion and expansion of a broad range of cell types, after functionalisation with protein nanosheets [24,29,30]. Upon assembly at Novec 7500/PBS interfaces, the interfacial shear storage modulus (iG’) of corresponding interfaces increased significantly and rapidly in all cases (Figure 3A). This was followed by the establishment of a plateau that persisted even following washing of the excess free protein from the aqueous phase. However, we noted an overall reduction in iG’ for interfaces stabilised by MA- and AC- BSA, compared to pristine BSA and NB-BSA (Figure 3A). This is perhaps reflecting the more significant reduction in the negative peaks seen in the CD spectra of the two former proteins (Figure 2B), implying a more prominent loss of α-helicity.

**Figure 3.**
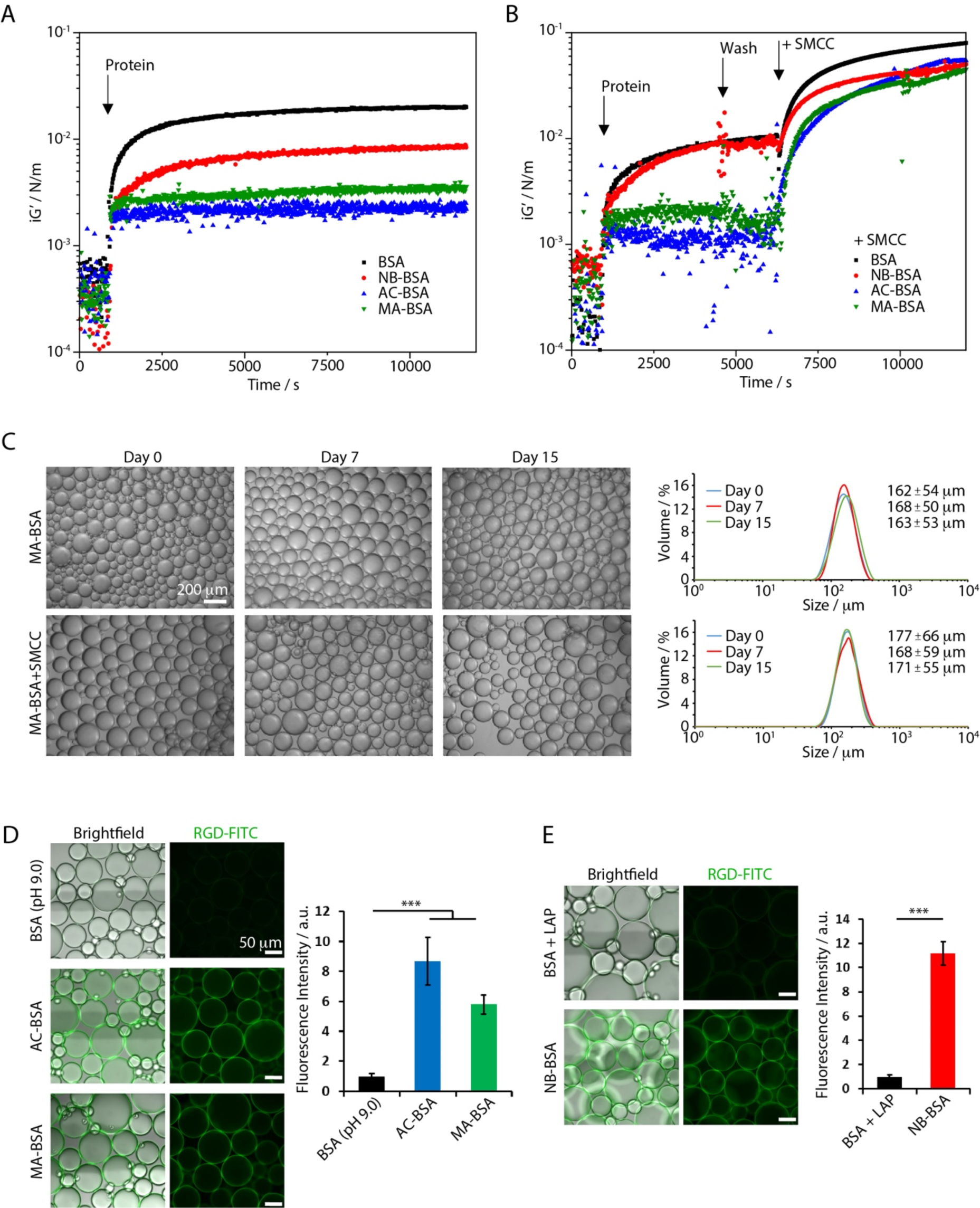
A. Evolution of the interfacial shear storage modulus (iG’) during the assembly of proteins at Novec 7500/PBS interfaces. BSA, NB-BSA, AC-BSA and MA-BSA at 1 mg/mL (0.1 Hz, 1.0 10^-4^ rad). B. Evolution of iG’ during BSA (1 mg/mL) assembly, followed by crosslinking with sulfo-SMCC at 2 mg/mL. C. Investigation of the stability of MA-BSA-based emulsions, for 15 days (microscopy images and volume of microdroplets evaluated via mastersizing), under agitation on an orbital shaker (60 rpm, 37°C). Values indicated are mean ± SD (N = 3). Scale bar, 200 μm. D. Bioemulsions formed after Michael addition reaction to conjugate CGGRGDSPK-FITC-G to BSA, AC-BSA and MA-BSA (brightfield/fluorescence microscopy images). Corresponding relative fluorescence intensity measured, to quantify bioconjugation (with respect to BSA-stabilised microdroplets). Values are mean ± SD (n > 20 droplets/condition; ***, P < 0.001). E. Bioemulsions formed after thiol-ene coupling to conjugate GCGGRGDSPK-FITC-G to BSA, and NB-BSA (brightfield/fluorescence microscopy images). Corresponding relative fluorescence intensity measured, to quantify bioconjugation (with respect to BSA-stabilised microdroplets). Values are mean ± SD (n > 20 droplets/condition; ***, P < 0.001). Scale bars, 50 μm.

As the interfacial shear moduli, elasticity and toughness of protein nanosheets was found to play a critical role in regulating cell adhesion and expansion at liquid-liquid interfaces [24,29–31], we investigated whether further crosslinking of the engineered proteins assembled could enhance corresponding interfacial mechanics. To do so, we used the heterobisfunctional crosslinker sulfo-SMCC, which was previously found to mediate the effective crosslinking of β-lactoglobulin at liquid interfaces [25,26]. Interfacial rheology was used to monitor changes in interfacial shear moduli during crosslinking (Figure 3B). To avoid protein aggregation in solution, we first washed the aqueous phase with PBS, prior to introduction of sulfo-SMCC. This led to a rapid increase in interfacial shear storage modulus for all three engineered proteins as well as pristine BSA, to 59-73 mN/m. Evaluation of different concentrations of sulfo-SMCC established that effective crosslinking was observed systematically above a concentration of 1 mg/mL (Supplementary Figure S1).

Examination of the frequency sweeps corresponding to the crosslinked and non-crosslinked interfaces further confirmed the striking increase in interfacial shear storage moduli in sulfo-SMCC treated assemblies (Supplementary Figure S2). In addition, whereas pristine BSA, and in particular engineered BSAs, displayed clear signs of failure at higher frequencies, all crosslinked nanosheets displayed stable viscoelastic profiles over the range of frequencies tested. Comparison of the interfacial shear storage (iG’) and loss moduli (iG’’) also revealed important differences in behaviour between pristine BSA and engineered albumins (Supplementary Figure S3). The iG’ of BSA was found to be higher than its iG’’, but the two moduli were closer in the case of NB-BSA and AC-BSA and iG’’ was higher than iG’ in the case of MA-BSA. This suggests significant levels of viscosity in engineered proteins. In contrast, all crosslinked nanosheets displayed significantly higher iG’, compared to their corresponding iG’’. To further confirm such findings, interfacial stress relaxation experiments were carried out (Supplementary Figure S4) [30]. These experiments demonstrated relatively elastic recovery profiles, with moderately high residual interfacial elastic stresses in the case of crosslinked nanosheets, whereas engineered albumin assemblies without sulfo-SMCC treatment displayed fast relaxation profiles typical of viscous materials.

Therefore, these results are consistent with the effective crosslinking, the strengthening and increase in elastic component of pristine BSA and engineered albumins treated with sulfo-SMCC. The reduction in interfacial shear moduli and the increase in viscous character of engineered protein assemblies in the absence of sulfo-SMCC, in particular for AC-BSA and MA-BSA, suggests that the partial denaturation of these proteins limits the establishment of protein networks with strong interfacial mechanics. This may indicate that protein denaturation is not only responsible to promote assembly at hydrophobic interfaces, but is also required for the physical crosslinking of associated networks. Such phenomena were found to be occurring more extensively in the case of pristine BSA and NB-BSA assemblies. However, variations in the surface densities of the various adsorbed proteins cannot be ruled out, although we note that the crosslinked nanosheets all displayed comparable interfacial shear moduli, suggestive of comparable surface densities.

The stability of emulsions formed with BSA and engineered albumins was next examined (Figure 3C and Supplementary Figure S5). The size of microdroplets was quantified by mastersizing and via brightfield microscopy. All emulsions, with and without treatment with sulfo-SMCC, were found to be stable over 15 days, at 37°C, in static conditions, as well as with agitation on an orbital shaker. Hence engineered albumins maintained excellent tensioactive properties for the stabilisation of emulsions, which were not significantly affected by crosslinking with sulfo-SMCC. Resulting bioemulsions were found to be more stable in culture conditions than previously reported poly(L-lysine) nanosheet-stabilised bioemulsions, which displayed modest signs of destabilisation (increase in droplet size) over 7 days of culture [33].

Finally, bioconjugation of the engineered proteins with cell adhesive peptides was achieved through Michael addition (for AC-BSA and MA-BSA) and radical thiol-ene coupling (for NB- BSA). Coupling-specific sequences, namely CGG-terminated for Michael addition, and GCGG-terminated for radical thiol-ene chemistry, were selected based on their established effectiveness to promote corresponding conjugations [50–54]. Fluorescence microscopy of resulting emulsions (after assembly of the protein bioconjugates), with FITC-tagged peptides, confirmed the specificity of these coupling strategies, compared to the pristine BSA control (Figures 3D and E). To further validate the coupling of peptides to NB-BSA via radical thiol-ene reaction we carried out gel electrophoresis, comparing native BSA, norbornylated BSA and RGD peptide functionalised NB-BSA (Supplementary Figure S6). Although the molecular weight of native BSA and functionalised BSA (e.g. NB-BSA) was found to be comparable, after peptide coupling the molecular weight of the resulting bioconjugate increased by approximately 10 kDa. Considering the molecular weight of the peptide selected (862 g/mol), coupling of approximately 12 peptide per protein was achieved, corresponding to 50% efficiency with respect to the number of norbornene residues per protein. Therefore, our results confirmed our ability to selectively bioconjugate engineered albumins enabling the stabilisation of bioemulsions, and further crosslinking with sulfo-SMCC for improved scaffolding properties.

The culture of MSCs on engineered protein nanosheet-stabilised bioemulsions was next investigated (Figure 4). Bone marrow-derived MSCs were cultured on RGD-coupled albumin nanosheet-stabilised bioemulsions, with and without SMCC crosslinking. Cell adhesion to uncrosslinked nanosheets was limited, with few clusters of cells seen at the surface of microdroplets 5 days post-seeding, but without evidence of extensive cell spreading or expansion (Figure 4A). Cells adhering to these uncrosslinked nanosheets did not assemble a structured cytoskeleton and did not form mature focal adhesions. In contrast, MSCs adhered and spread extensively at the surface of crosslinked nanosheets and corresponding bioemulsion microdroplets (Figure 4A). After 5 days of culture, these cells displayed a well-structured cytoskeleton, with apparent mature stress fibres radiating from focal adhesions. Hence the adhesive machinery, underpinned by actomyosin assembly and contractility, and mediated by integrin clustering promoting focal adhesion development was found to be engaged in adherent cells spreading at the surface of bioemulsions. Considering the role of this machinery in the maintenance of stem cell phenotype and fate decision [4,55,56], these observations suggested SMCC-crosslinked bioemulsions could sustain the expansion of bone marrow derived MSCs.

**Figure 4.**
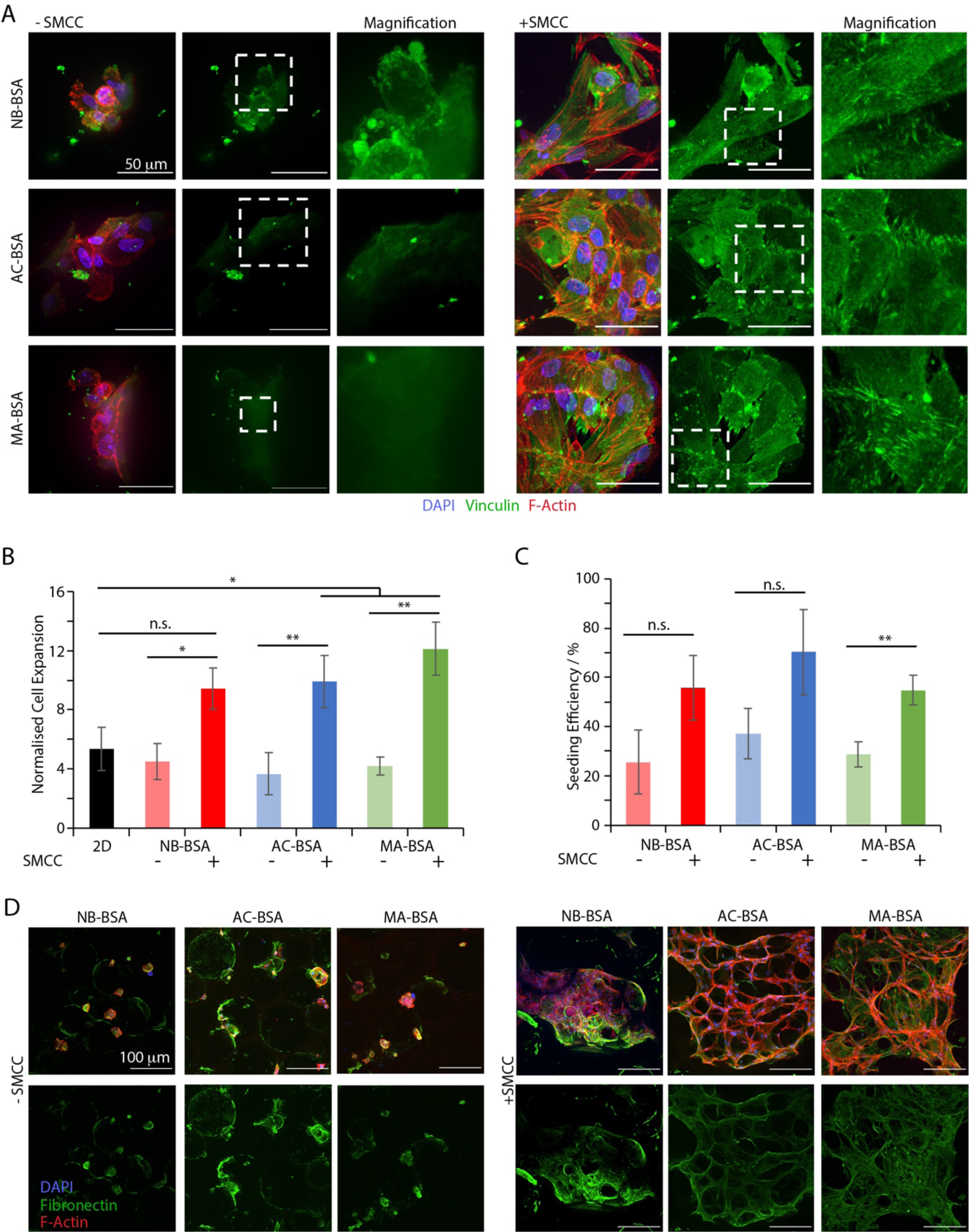
A. Immunofluorescence microscopy images of focal adhesions (vinculin, green) formed by MSCs adhering to functionalised and crosslinked BSA-stabilised bioemulsions for 5 days. Magnifications show mature FAs on crosslinked interfaces but not on interfaces untreated with sulfo-SMCC. B. Quantification of MSC expansion for 7 days of culture in 2D and at the surface of crosslinked and non-crosslinked NB-BSA, AC-BSA and MA-BSA nanosheets stabilising corresponding bioemulsions. Quantification via CyQuant assay (respective to cell numbers at day 1). Values are mean ± SD of from 3 different experiments (n.s., non-significant, P > 0.05; *, P< 0.05; **, P < 0.01). C. Corresponding initial seeding efficiencies, characterised by CyQuant assay (with respect to initial number of cells seeded). Values are mean ± SD from 3 different experiments (n.s., non significant, P > 0.05; **, P < 0.01). D. Matrix deposition (fibronectin, green) on bioconjugated BSA-stabilised bioemulsions, with or without sulfo-SMCC crosslinking, after culture of MSCs for 7 days. Note that crosslinking of bioconjugated-BSA nanosheets promoted the assembly of an ECM network (fibronectin) at corresponding interfaces, extending into a 3D matrix.

In turn, cell expansion and initial seeding efficiency were also found to be enhanced on bioemulsions crosslinked with SMCC (Figures 4B and C). Significant increases in cell densities were observed on all crosslinked bioemulsions, with densities overtaking those observed even on 2D plastic cultures, in the case of MA-BSA. In agreement with the high spreading observed at day 5, MSCs cultured on crosslinked nanosheets adhered more effectively (high initial seeding efficiencies achieved) and proliferated more rapidly than cells adhering to uncrosslinked counterparts. Considering the importance of ECM ligation and matrix mechanics in supporting cell adhesion, these results indicate that the important change in nanoscale mechanics achieved through the crosslinking of nanosheets is essential to promote MSC expansion on bioemulsions. This result is in agreement with observations made with poly(L-lysine) nanosheets that had been found to sustain stem cell adhesion and expansion when displaying sufficiently strong and elastic interfacial mechanics [24,30,31].

MSCs often play an important role in assembling, remodelling and maintaining ECM in corresponding niches, for example in the bone marrow [57–59]. Examination of matrix deposition after 7 days of culture on bioemulsions indeed confirmed the deposition of substantial levels of fibronectin (Figure 4D). This process was again modulated by interfacial mechanics and nanosheet crosslinking, as uncrosslinked bioemulsions supported only the deposition of patchy fibronectin domains, potentially associated with the poor adhesion and expansion observed at these interfaces. In contrast, significant fibronectin deposition, extending between droplets aggregating together into small tissue-like structures were observed on SMCC-crosslinked bioemulsions. This was particularly striking in the case of AC- BSA and MA-BSA, perhaps suggesting that residual acrylate and methacrylate moieties may contribute to such matrix deposition, whereas norbornyl residues do not. Indeed, thiols and amines may bind to acrylates and methacrylates in physiological conditions, via Michael addition, whereas norbornyl groups are unable to sustain coupling without activation (e.g. through radical initiation). Therefore MSCs growing on bioemulsions are able to assemble and remodel their extra-cellular microenvironment, recreating tissue-like structures resembling adipose tissues. Such phenomena is attractive to the formation of stem cell niches in vitro, for example for the culture of hematopoietic stem cells [34].

To demonstrate the scalability of bioemulsions for MSC culture, we next selected MA-BSA stabilised microdroplets for culture in conical flask bioreactors. The selection of 250 mL conical flasks with vented caps (Figure 5A) allowed us to scale up MSC culture 100 times compared to culture on multi-well plates. Hence 3 M cells were seeded on RGD-coupled MA- BSA bioemulsions and cultured for 15 days in a conical flask bioreactor, on an orbital shaker. Although seeding efficiencies were found to decrease compared to those achieved in 2D cultures, they remained close to 60% (Figure 5B), comparing favourably with seeding protocols on solid microcarriers [60,61]. Indeed, seeding protocols on microcarriers face the challenge of promoting cell adhesion sufficiently fast to compete with the sedimentation of cultures. The high seeding densities achieved enabled the sustaining of homogenous cell coverage and expansion in the following 15 days, with bioemulsion cultures displaying high confluency (Figure 5C). Throughout this culture time, cell viability was found to remain unchanged (typically above 95%) and comparable for cells cultured on bioemulsion and on 2D tissue culture plastic (Figures 5D and E).

**Figure 5.**
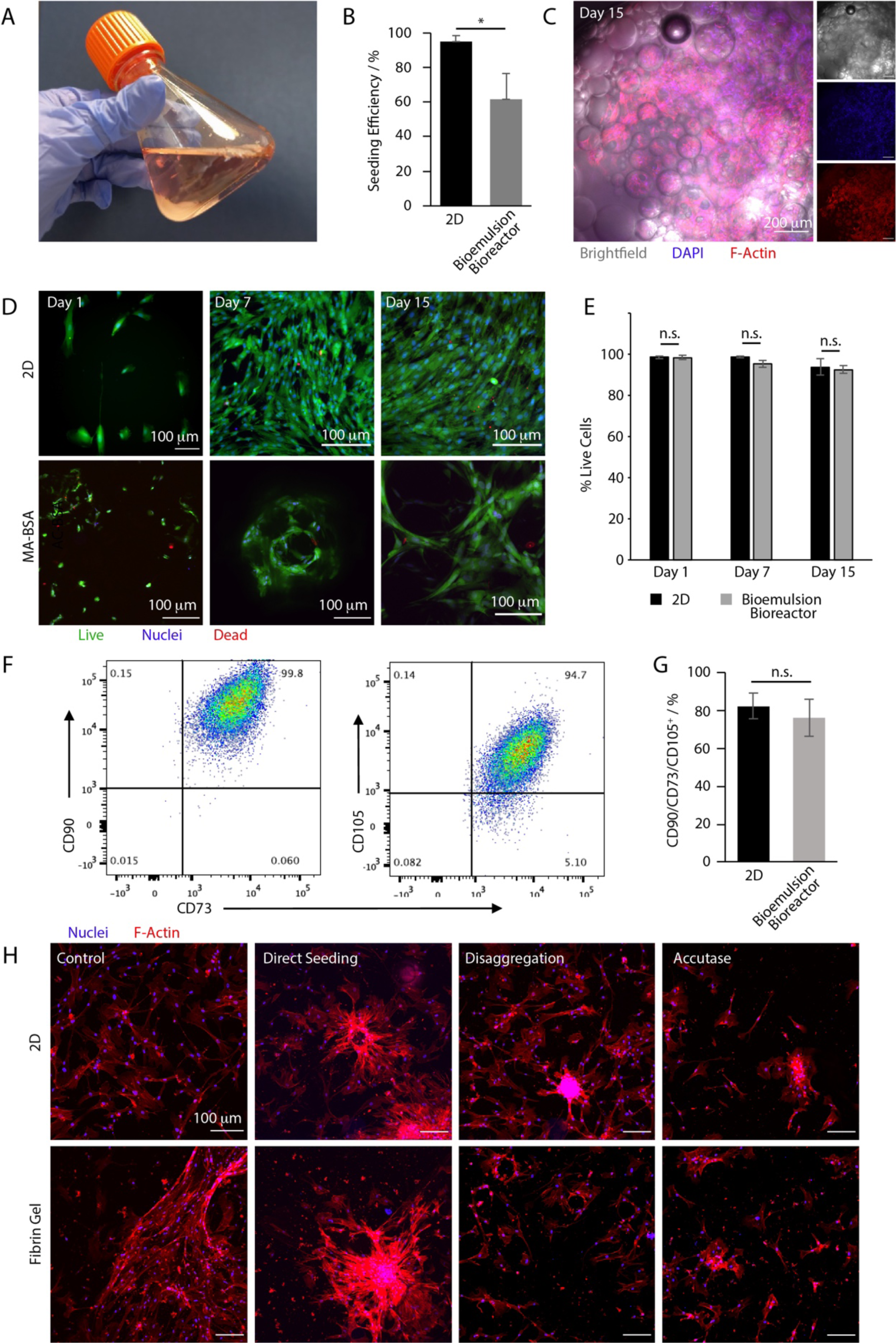
Scale up of MSC culture on bioemulsions. A. Image of cell culture conical flask used to scale up MSC culture 100 fold. MA-BSA stabilised bioemulsion (SMCC crosslinked and RGD conjugated) B. Seeding efficiency achieved on bioemulsion, characterised by CyQuant assay (with respect to initial number of cells seeded). Values are mean ± SD from 3 different experiments (*, P < 0.05). C. Brightfield/fluorescence microscopy images of MSCs grown on emulsion for 15 days. Scale bar, 200 μm. D. Cell viability assessed at days 1, 7 and 15 (live/dead assay), on 2D culture plastic and MA-BSA stabilised bioemulsion (SMCC crosslinked and RGD conjugated). E. Corresponding quantification of data. Values are mean ± SD from 3 different experiments (n.s., P > 0.05). F. Characterisation of CD73, CD90 and CD105 expression in MSCs cultured on MA-BSA stabilised bioemulsions (SMCC crosslinked and RGD conjugated) for 15 days (FACS data). G. Corresponding quantification of triple marker expression for cells grown on 2D culture plastic and bioemulsion. Values are mean ± SD from 3 different experiments (n.s., P > 0.05). H. Fluorescence microscopy images of MSCs cultured on bioemulsion for 15 days and then seeded on 2D culture plastic or in fibrin gels for 5 days, following different processing protocols: Control, MSCs cultured on 2D tissue culture plastic prior to reseeding; Direct seeding, cell-seeded droplets directly deposited or embedded; Disaggregation, emulsion cultures passed through a thin needle to induce cell detachment; Accutase, cell detachment using Accutase, prior to seeding on 2D or in fibrin gels.

The phenotype of MSCs cultured on bioemulsions was examined next. Specifically, we quantified the co-expression of the membrane markers CD73, CD90 and CD105, three positive markers of bona fide MSCs [62], after 15 days of culture, via FACS (Figure 5F). Overall, >99% cells remained CD73+ and CD90+, and 95% cells were CD105+, after 15 days culture on MA- BSA emulsions, with overall triple positive populations equivalent to those of MSCs cultured on 2D tissue culture plastic (Figure 5G). Triple marker expression typically correlates tightly with the capacity of MSCs to differentiate in multiple lineages [33,60,62] and is considered to be a good predictor of therapeutic performance [59]. Therefore, key features defining MSC phenotype and therapeutic capacity were maintained on MA-BSA bioemulsions.

The ability to recover cells rapidly, whilst limiting processing steps and potential cell loss, is often considered an important hurdle to the scale up of cell manufacturing. We explored two methods of cell recovery requiring no-to-limited processing, compared to enzymatic cell detachment (Figure 5H). Cell colonies growing around droplets were either directly deposited on 2D tissue culture plastic and allowed to adhere and spread, transferring to this new substrate, or directly embedded in fibrin gel, to promote cell migration in a 3D matrix. Alternatively, cells adhering to emulsions were gently dissociated by pipetting through narrow pipettes, prior to transfer to 2D substrates or fibrin gels. Both methods enabled cell transfer and recovery, similarly to what was observed for cells recovered via enzymatic treatment, from bioemulsions or after culture on 2D substrates. Therefore, bioemulsions also enable simpler cell recovery, without enzymatic treatment, reducing processing time and costs and potentially limiting cell losses.

Direct implantation of cells following culture is attractive as limiting processing steps, time and costs, but the implantation of fluorinated oils, even with good cytocompatibility profile and inertness, as for Novec 7500, pauses hurdles to regulatory approval. Instead, other oil types such as silicone, mineral and plant-based oils constitute more attractive candidates for translation, and to allow direct implantation of cell colonies growing around droplets. To demonstrate the versatility of the engineered proteins developed for bioemulsion design, the culture of MSCs on plant-based oil emulsion (rapeseed oil) stabilised by NB-BSA, AC-BSA and MA-BSA was investigated. After 7 days of cultures, MSCs could be seen spreading around rapeseed oil microdroplets coated by all three proteins and crosslinked with SMCC (Figure 6A). Cell densities steadily increased at each time points for all three protein-stabilised bioemulsions, although this increase was more pronounced in the case of MA-BSA (Figure 6B).

**Figure 6.**
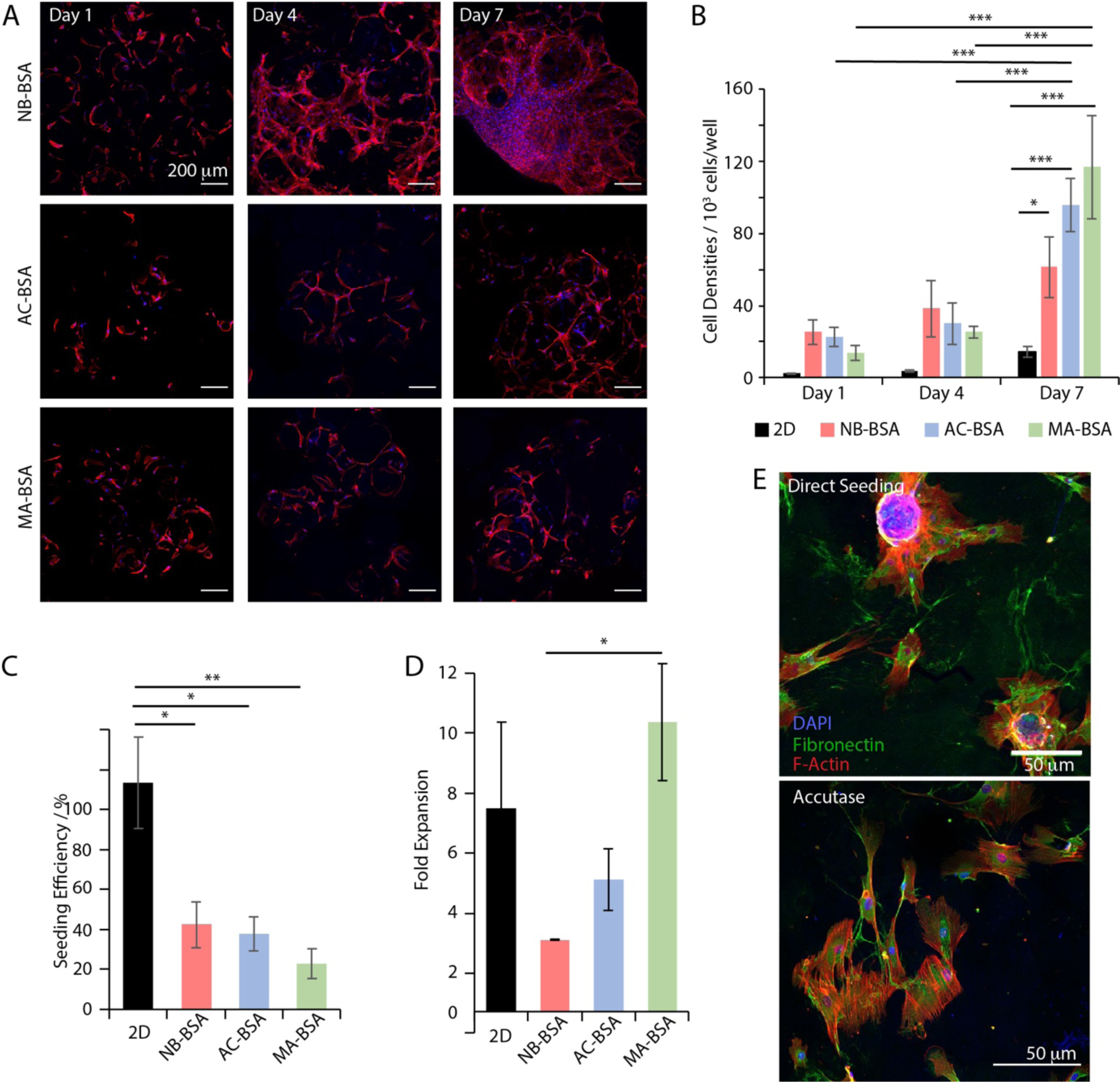
A. Fluorescence microscopy images of MSCs cultured on rapeseed oil bioemulsions stabilised by NB-BSA, AC-BSA and MA-BSA, crosslinked with SMCC, at different timepoints. B. Quantification of cell densities of MSCs cultured on 2D plastic and on bioemulsions, at different timepoints, using a CyQuant assay. Values are mean ± SD from 3 different experiments (*, P < 0.05; ***, P < 0.001; ****, P< 0.0001). C and D. Corresponding seeding efficiencies and fold expansion, respectively. Values are mean ± SD from 3 different experiments (*, P < 0.05; **, P < 0.01; ***, P < 0.001). E. MSCs grown on 2D tissue culture plastic for 2 days after harvesting from a MA-BSA bioemulsion culture (at day 7). Re-seeding: direct reseeding of cell-seeded microdroplets; trypsin, enzymatic cell detachment prior to seeding.

Although seeding efficiencies were lower than those determined for fluorinated oil emulsions (Figure 6C, compared to Figure 5B), cell densities caught up rapidly with those measured for 2D cultures and, in the case of MA-BSA stabilised bioemulsions, expansion folds reached comparable or slightly higher levels (Figure 6D). The reduced seeding efficiency is likely due to the buoyancy of plant-based bioemulsions. Indeed, with these bioemulsions, care had to be taken whilst depositing the cell suspension, so that cells could gently sediment whilst passing through the emulsion layer, to enable adhesion. Once cells had sedimented below this emulsion phase, they had no further opportunity to adhere to adhesive microdroplets and therefore this resulted in limited seeding efficiencies. However, this process could be potentially further optimised by increasing the height of the emulsion phase, or increasing the viscosity of the loading medium. Finally, MSCs cultured on rapeseed oil bioemulsions were reseeded, either directly after gentle disaggregation, or after enzymatic treatment (Figure 6E), displaying capacity to re-adhere to a new substrate and comparable potential for the simplification of cell processing to Novec 7500 bioemulsions.

## Discussion and conclusions

A growing number of reports has demonstrated the importance of local physical and mechanical properties on the regulation of cell adhesion and phenotype [63–66]. The rational design of liquid-liquid interfaces to promote cell adhesion, expansion and stemness retention is in excellent agreement with this literature and illustrates the importance of local nanoscale mechanics in regulating cell phenotype. This concept offers an unprecedented flexibility of engineering for the design of novel biomaterials for application in the fields of biomanufacturing, regenerative medicine and tissue engineering. As bioconjugation tools and the control of tensioactive properties are well-established, an important component of design to enable cell culture at liquid interfaces is the control of interfacial shear mechanics. These properties, rather than dilatational mechanics, underpin the design of protein nanosheets, distinguishing these interfaces from more classic protein assemblies. In this context, sulfo-SMCC is one example of crosslinker that demonstrated consistent performance for the various albumins explored in this study. However, its precise crosslinking mechanism remains only partly established. Indeed it could originate from hetero-bicoupling within protein assemblies, but could equally arise from hydrophobic interactions, which have been proposed to dominate the crosslinking of protein nanosheets assembled with hydrophobic co-surfactants [30,31].

The combination of these three properties for the rationale design of interfaces enables the development of bioactive emulsions. Bioemulsion design can increasingly make use of materials approved by regulators for a range of procedures and permanent implantation (e.g. oils and proteins of interest), without relying on cytotoxic components. Costs are also reduced through such design, broadening their scope of applications. The use of engineered proteins, which could be produced recombinantly, and the possibility of using rapeseed oil and other plant-based oils as substrates are important steps towards further translation of bioemulsions. In this context, engineered globular proteins are well-suited to stabilise a broad range of such oils, extending on concepts developed for formulation in the food industry.

Some hurdles remain to be addressed for translation, for example allowing the improvement of initial cell seeding. This can however be addressed through a combination of emulsion engineering (e.g. regulating size distribution and viscosity), as well as process engineering (e.g. the design of devices enabling to maximise cell adhesion during sedimentation, in a scalable flowable process. The advances in bioemulsion design achieved may allow tackling other pitfalls in cell manufacturing, bypassing the complex processing steps typically required to recover cell masses, e.g. through centrifugation, direct cell disaggregation, or altogether bypassing any processing required, allowing direct implantation following culture. These advantages may find application in other fields, such as in cultivated meat, where cell output, processing and costs are critical hurdles to the translation of tissue culture technologies to the development of affordable and scalable food products. Overall, bioemulsions present unique opportunities to transform biomanufacturing and bioprocessing for cell therapies, not only enabling scale up in safe and cost-effective formats, but also allowing to re-think harvesting and implantation protocols and strategies to speed up the translation of novel cell technologies.

## Acknowledgement

We thank the European Research Council (ProLiCell, 772462) for support.

## Conflict of Interest

The authors declare no conflict of interest.

## Notes

### Competing Interest Statement

The authors have declared no competing interest.

