## Supplementary Materials for "Engineered Protein Nanosheets for the Scale up of Mesenchymal Stem Cell Culture on Bioemulsions"

### Supplementary Information

Minerva Bosch-Fortea[1], Clemence Nadal[1], Alexandra Chrysanthou[1], Daniele Marciano[1], Hassan Kanso[1], Nam Nguyen[1] and Julien E. Gautrot[1*]

[1] School of Engineering and Materials Science, Queen Mary University of London, Mile End Road, London E1 4NS, United Kingdom.

* Correspondence:

Julien E. Gautrot

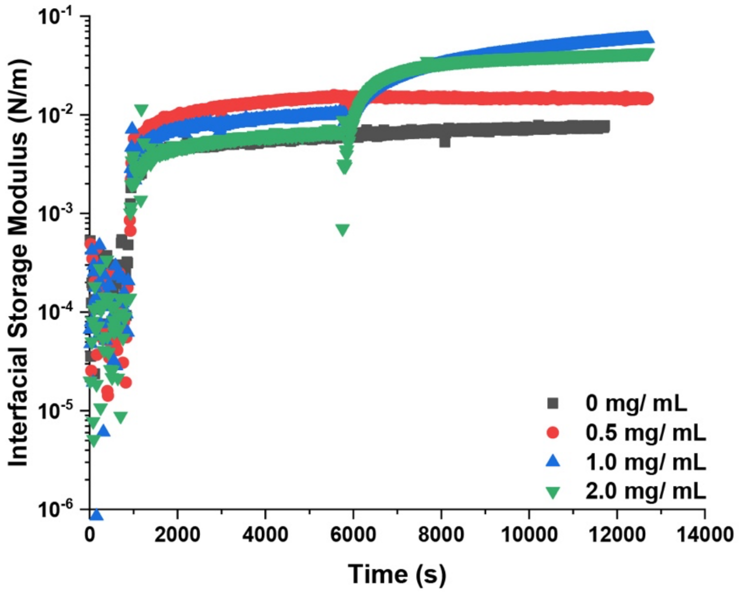


**Supplementary Figure S1.** Evolution of the interfacial shear storage modulus (iG') during the assembly of NB-BSA (1 mg/mL) at Novec 7500/PBS interfaces, followed by crosslinking with sulfo-SMCC at different concentrations (0.1 Hz, 1.0 $10^{-3}$ rad).

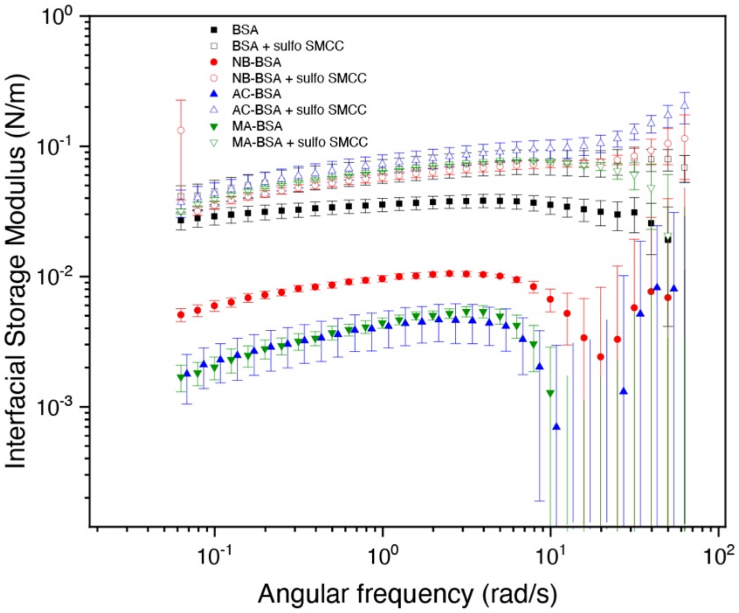


**Supplementary Figure S2.** Frequency sweep profiles recorded at a displacement of 1.0 x $10^{-3}$ rad for the different protein assemblies with and without crosslinking with sulfo-SMCC. Error bars are s.e.m.; n = 3.

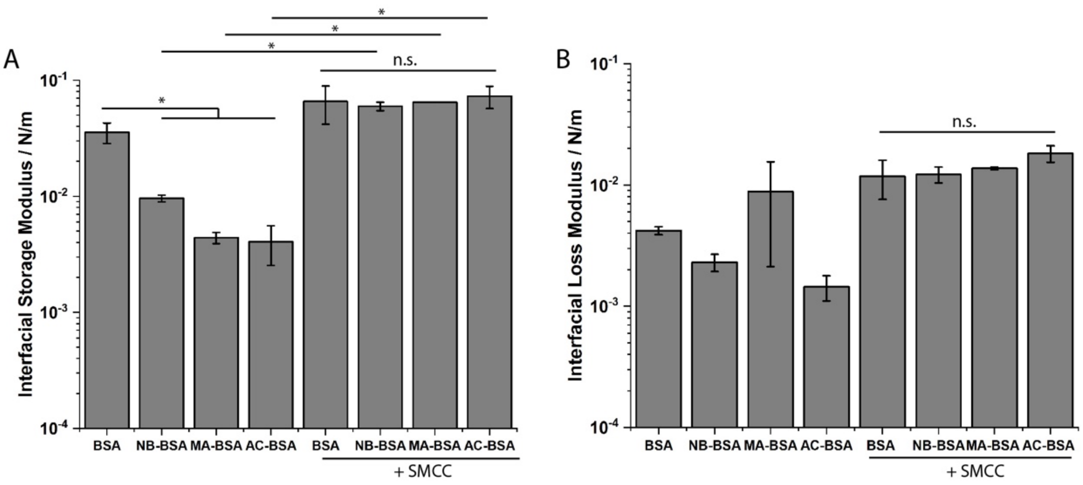


**Supplementary Figure S3.** Interfacial storage and loss moduli measured at 0.1 Hz in the frequency sweeps (error bars are s.e.m.; n = 3).

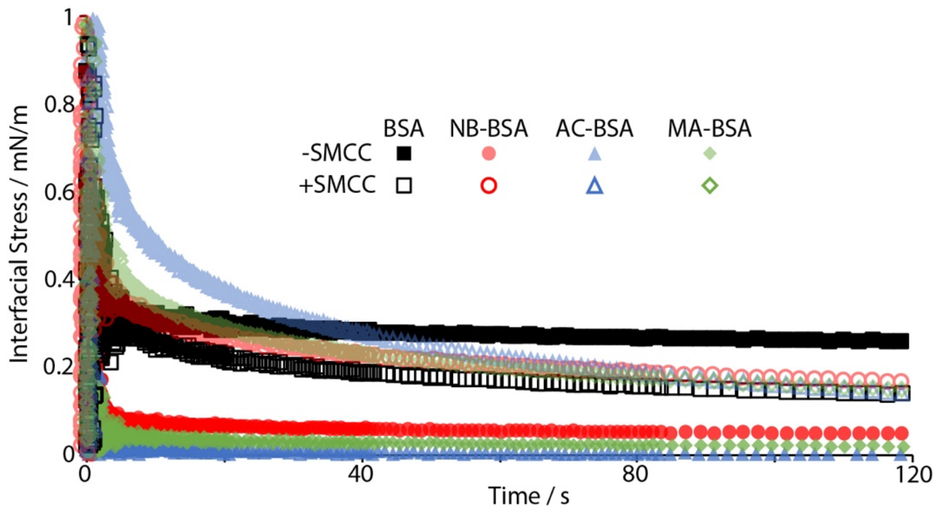


**Supplementary Figure S4.** Viscoelasticity of engineered albumins nanosheets formed at PBS/Novec oil interfaces: normalised stress relaxation profiles for the different proteins with and without crosslinking with sulfo-SMCC, recorded after 0.5 % strain.

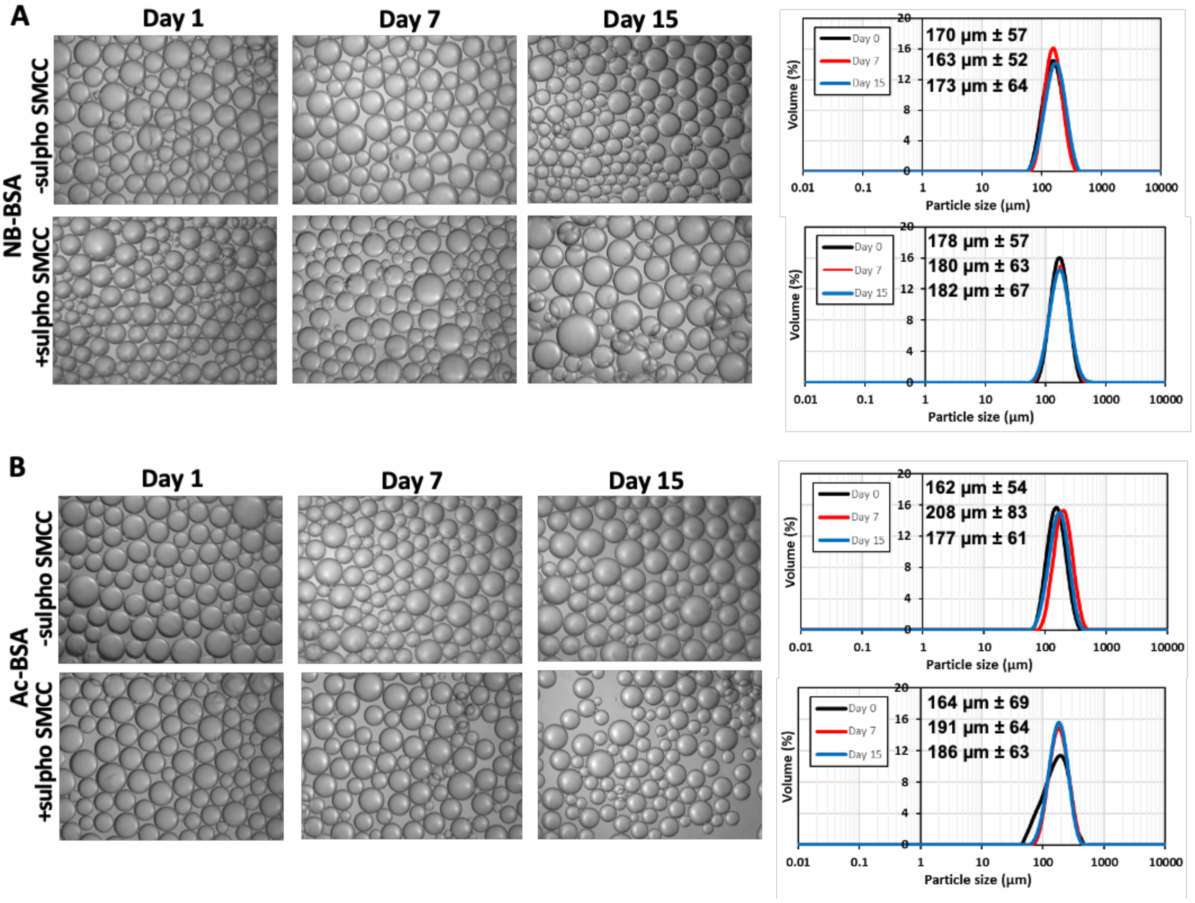


**Supplementary Figure S5.** BSA-based nanosheets allow the formation bioemulsions that remain stable under agitation over 15 days. The volume of fluorinated oil bioemulsions was evaluated over time using mastersizing. NB-BSA (A), AC-BSA (B) stabilised microdroplets for > 15 days under agitation (orbital shaker, 60 rpm, 37$^{o}$C). Brightfield images qualitatively confirmed the maintenance of droplet sizes over time. Scale bar, 200 µm. Graphs show particle size distribution at days 0, 7 and 15. Values are mean ± SD from triplicate measurements.

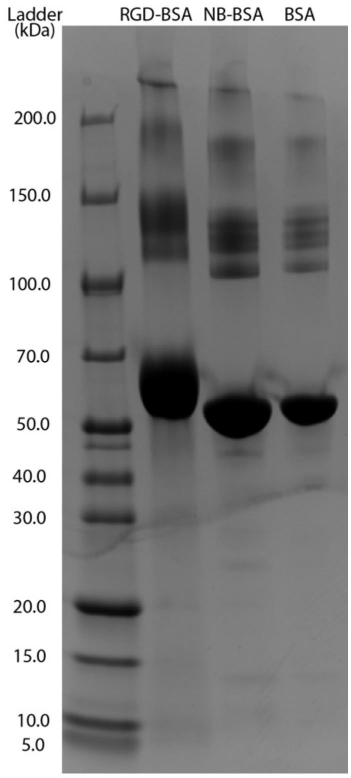


**Supplementary Figure S6.** Gel electrophoresis confirms the coupling of GCGGRGDSPG sequences to NB-BSA.

**Supplementary Tables**

**Supplementary Table S1** Summary of statistical analysis of the data presented in Figure 3D.

| FIGURE 3D | Probability | Significance |
|---|---|---|
| BSA vs. AC-BSA | 2.22E-15 | *** |
| BSA vs. MA-BSA | 1.28E-16 | *** |

**Supplementary Table S2** Summary of statistical analysis of the data presented in Figure 3E.

| FIGURE 3E | Probability | Significance |
|---|---|---|
| BSA vs. NB-BSA | 7.42E-17 | *** |

**Supplementary Table S3** Summary of statistical analysis of the data presented in Figure 4B.

| FIGURE 4B | Probability | Significance |
|---|---|---|
| 2D vs. NB-BSA | 0.9956 | n.s |
| 2D vs. NB-BSA + SH-SMCC | 0.138 | n.s |
| 2D vs. AC-BSA | 0.8903 | n.s |
| 2D vs. AC-BSA + SH-SMCC | 0.0795 | n.s |
| 2D vs. MA-BSA | 0.9781 | n.s |
| 2D vs. MA-BSA + SH-SMCC | 0.0049 | ** |
| NB-BSA vs. NB-BSA + SH-SMCC | 0.0487 | * |
| NB-BSA vs. AC-BSA | 0.9966 | n.s |
| NB-BSA vs. AC-BSA + SH-SMCC | 0.0271 | * |
| NB-BSA vs. MA-BSA | 1 | n.s |
| NB-BSA vs. MA-BSA + SH-SMCC | 0.0017 | ** |
| NB-BSA vs. SH-SMCC vs. AC-BSA | 0.0172 | * |
| NB-BSA+SH-SMCC vs. AC-BSA + SH-SMCC | 0.9999 | n.s |
| NB-BSA + SH-SMCC vs. MA-BSA | 0.0328 | * |
| NB-BSA + SH-SMCC vs. MA-BSA + SH-SMCC | 0.5263 | n.s |
| AC-BSA vs. AC-BSA + SH-SMCC | 0.0095 | ** |
| AC-BSA vs. MA-BSA | 0.9998 | n.s |
| AC-BSA vs. MA-BSA + SH-SMCC | 0.0006 | *** |
| AC-BSA+SH-SMCC vs. MA-BSA | 0.0182 | * |
| AC-BSA + SH-SMCC vs. MA-BSA + SH-SMCC | 0.7149 | n.s |
| MA-BSA vs. MA-BSA + SH-SMCC | 0.0011 | ** |

**Supplementary Table S4** Summary of statistical analysis of the data presented in Figure 4C.

| FIGURE 4C | Probability | Significance |
|---|---|---|
| NB-BSA vs. NB-BSA + SH-SMCC | 0.0814731 | n.s |
| AC-BSA vs. AC-BSA + SH-SMCC | 0.07963902 | n.s |
| MA-BSA vs. MA-BSA + SH-SMCC | 0.00948041 | ** |

**Supplementary Table S5** Summary of statistical analysis of the data presented in Figure 5B.

| FIGURE 5B | Probability | Significance |
|---|---|---|
| 2D vs. Bioemulsion Bioreactor | 0.04042876 | * |

**Supplementary Table S6** Summary of statistical analysis of the data presented in Figure 5E.

| FIGURE 5E | Probability | Significance |
|---|---|---|
| Day 1 2D vs. Bioemulsion Bioreactor | 0.87673473 | n.s |
| Day 7 2D vs. Bioemulsion Bioreactor | 0.13301768 | n.s |
| Day 14 2D vs. Bioemulsion Bioreactor | 0.58697341 | n.s |

**Supplementary Table S7** Summary of statistical analysis of the data presented in Figure 5G.

| FIGURE 5G | Probability | Significance |
|---|---|---|
| | 0.44468891 | n.s |

**Supplementary Table S8.** Summary of statistical analysis of the data presented in Figure 6B.

| FIGURE 6B | Probability | Significance |
|---|---|---|
| NB & day 1 vs. NB & day 4 | 0.994 | n.s |
| NB & day 1 vs. NB & day 7 | 0.1931 | n.s |
| NB & day 4 vs. NB & day 7 | 0.765 | n.s |
| AC & day 1 vs. AC & day 4 | 1 | n.s |
| AC & day 1 vs. AC & day 7 | 0.0002 | *** |
| AC & day 4 vs. AC & day 7 | 0.0008 | *** |
| MA & day 1 vs. MA & day 4 | 0.9977 | n.s |
| MA & day 1 vs. MA & day 7 | 0.00000072 | **** |
| MA & day 4 vs. MA & day 7 | 0.00000571 | **** |
| 2D & day 1 vs. 2D & day 4 | 1 | n.s |
| 2D & day 1 vs. 2D & day 7 | 0.9976 | n.s |
| 2D & day 4 vs. 2D & day 7 | 0.999 | n.s |
| NB & day 1 vs. AC & day 1 | 1 | n.s |
| NB & day 1 vs. MA & day 1 | 0.9974 | n.s |
| NB & day 1 vs. 2D & day 1 | 0.7684 | n.s |
| AC & day 1 vs. MA & day 1 | 0.9998 | n.s |
| AC & day 1 vs. 2D & day 1 | 0.8799 | n.s |
| MA & day 1 vs. 2D & day 1 | 0.9984 | n.s |
| NB & day 4 vs. AC & day 4 | 0.9999 | n.s |
| NB & day 4 vs. MA & day 4 | 0.9934 | n.s |
| NB & day 4 vs. 2D & day 4 | 0.2274 | n.s |
| AC & day 4 vs. MA & day 4 | 1 | n.s |
| AC & day 4 vs. 2D & day 4 | 0.585 | n.s |
| MA & day 4 vs. 2D & day 4 | 0.8214 | n.s |
| NB & day 7 vs. AC & day 7 | 0.2342 | n.s |
| NB & day 7 vs. MA & day 7 | 0.006 | ** |
| NB & day 7 vs. 2D & day 7 | 0.0296 | * |
| AC & day 7 vs. MA & day 7 | 0.8428 | n.s |
| AC & day 7 vs. 2D & day 7 | 0.00003708 | **** |
| MA & day 7 vs. 2D & day 7 | 0.00000079 | **** |

**Supplementary Table S9.** Summary of statistical analysis of the data presented in Figure 6C.

| FIGURE 6C | Probability | Significance |
|---|---|---|
| 2D vs. NB-BSA | 0.0174959 | * |
| 2D vs. AC-BSA | 0.01212562 | * |
| 2D vs. MA-BSA | 0.00608677 | ** |

**Supplementary Table S10.** Summary of statistical analysis of the data presented in Figure 6D.

| FIGURE 6D | Probability | Significance |
|---|---|---|
| 2D vs. NB-BSA | 0.09359006 | n.s |
| 2D vs. AC-BSA | 0.32408954 | n.s |
| 2D vs. MA-BSA | 0.09359006 | n.s |
| NB-BSA vs. AC-BSA | 0.05270984 | n.s |
| NB-BSA vs. MA-BSA | 0.03760574 | * |
| AC vs. BSA-MA vs. BSA | 0.11870761 | n.s |
